# Biomimetic porous silicon for rationally engineered combination therapy to induce immunogenic cell death

**DOI:** 10.1101/2024.11.28.625849

**Authors:** Silja Saarela, Emilia Happonen, Milla-Iida Laari, Zakir Mohammed, Shaghayegh Saba, Ruixian Liu, Juha Klefström, Vesa-Pekka Lehto, Wujun Xu

## Abstract

Combined chemo-photothermal therapy (CHT-PTT) is a promising treatment for metastatic cancers. It enables the synergistic induction of immunogenic cell death (ICD), while sensitising suppressive tumour microenvironment to immunotherapy. To induce ICD, the rational design of a combined CHT-PTT regimen remains unclear. In the present study, black porous silicon nanoparticles (BPSi NPs) were utilized as both drug carriers and photothermal conversion agents. To enhance their colloidal stability and homotypic targeting, BPSi NP surfaces were modified with polyethylene glycol and further coated with cancer cell membranes (CMs). Six chemotherapeutic drugs with different cell growth inhibition mechanisms were tested against metastatic MDA-MB-231 breast cancer cells to evaluate their ICD induction potentials. Finally, in the combined CHT-PTT, the effect of temperature was evaluated. Based on the analysis of ICD markers, high-mobility group box 1 and calreticulin proteins, we found that bromodomain-containing protein 4 inhibitor, (+)-JQ-1 (JQ1) was the optimal drug, and the mild-hyperthermia temperature of 45 °C was the best temperature setting in the combined CHT-PTT to induce ICD. Thus, our study provides rationally designed CHT-PTT regimen for efficient cancer treatment with high efficacy and low side effects.

## 1. Introduction

Of breast cancers, 15 % are classified as metastatic triple negative breast cancers (TNBCs) (1). TNBC involves a lack of oestrogen and progesterone receptors and human epidermal growth factor receptor 2 (HER2), which have a major role as molecular targets of therapeutic agents. Thus, the ineffectiveness of targeted therapies leads to poor prognosis and overall survival (1,2). TNBC is commonly treated by conventional chemotherapy and radiotherapy to inhibit the primary tumour growth and spread (3). However, these treatment methods involve severe side effects and tumour recurrence emphasizing the importance of new therapies (1,3). One promising approach for treating metastatic cancers is immunotherapy. Despite of the potential, its immunotherapeutic efficacy is limited because of the suppressed tumour microenvironment of most tumours. One possible solution to sensitize the tumour microenvironment to immunotherapy is via the induction of immunogenic cell death (ICD) (4). ICD is a type of programmed cell death that enhances tumour antigen occurrence (5). It can be achieved by releasing soluble damage-associated molecular patterns (DAMPs), such as calreticulin (CRT) and high-mobility group box 1 (HMGB1) proteins (6). As cancer cells undergo ICD, CRT translocates from the endoplasmic reticulum (ER) to concentrate on the cell surface, acting as an “eat me “ signal (4,7). Additionally, HMGB1 is released from the cell nucleus to the extracellular space (6,7). The function of these ICD markers is to activate immune cells (7,8) and convert tumour microenvironments from being immunologically “cold” to “hot” (4).

ICD can be induced by chemotherapy (CHT) and physical modalities such as photothermal therapy (PTT) (9). Some chemotherapeutic drugs, by inducing ICD, can facilitate immune-mediated recognition of cancer cells (5). However, chemotherapeutic drugs have a tendency to cause several side effects and lead to chemoresistance, rendering regular CHT for ICD induction unfeasible (10,11). For targeted cancer cell death, PTT is a non-invasive and local treatment method (12,13). It typically uses 650-900 nm near-infrared (NIR) light that is converted to heat, causing hyperthermia in the tumour tissue via nanoparticles (NPs) as photothermal conversion agents (PCAs) (6,12). Inducing ICD purely with PTT is challenging, because its treatment volumes and durations pose limitations (13), and laser power causes partial heat ablation in tumours (14). In addition, for disseminated tumours, metastasis occurs in regions unreachable by NIR light, rendering PTT impractical (15). Individual challenges in CHT and PTT inspired us to combine the best features of both methods to synergistically induce ICD by NP-based combined chemo-photothermal therapy (CHT-PTT). CHT-PTT enables a lower drug dose, reducing side effects and chemoresistance (9). Moreover, by improving ICD induction efficiency, CHT-PTT could prevent tumour recurrence and inhibit tumour metastasis (16).

NPs possess multifunctional properties, rendering them as a great platform for CHT-PTT (14). Especially suitable are black porous silicon (BPSi) NPs, acting simultaneously as drug carriers and PCAs (4). BPSi NPs are biocompatible and possess a large surface area and pore volume (12,17), permitting high drug-loading capacity and sustained drug release (10,18). Because of their highly light-absorbing black colour, BPSi NPs have high photothermal conversion efficiency (33.4 %), rendering them ideal PCAs for PTT (12). BPSi NP surfaces are easily modified, enabling further functionalization for drug absorption and release (12,19). For surface functionalization, to increase NP colloidal stability, a well-known method is polyethylene glycol (PEG) coating (20,21). A new strategy to improve NP colloidal stability and homologous targeting is coating with cancer cell membranes (CMs) (22,23). CM coating enhances biocompatibility, cellular uptake, and sustained drug release, rendering these biomimetic NPs a notable option for CHT-PTT to induce ICD (23,24). Here, we applied BPSi NPs in CHT-PTT as drug carriers and PCAs to synergistically induce ICD in MDA-MB-231 cells (TNBC cell line) expressing HMGB1 green fluorescence protein (HMGB1-GFP) and CRT red fluorescence protein (CRT-RFP), *in vitro*. Briefly, to enhance their biocompatibility and colloidal stability, BPSi NPs were PEGylated and CM-coated. Homotypic targeting properties of CM-PEG-BPSi NPs were confirmed via confocal laser scanning microscopy (CLSM) imaging. Further, we selected six chemotherapeutic drugs with different cell growth inhibition mechanisms. These six were mitoxantrone (MTX), doxorubicin (DOX), paclitaxel (PTX), dasatinib (DAS), omipalisib (GSK), and (+)-JQ-1 (JQ1) (Figure 1). To study the potential of these drugs for ICD induction in temperature-controlled PTT, biomimetic NPs were combined with their subtherapeutic doses. To test the optimal temperature to induce ICD in CHT-PTT, based on these free drug experiments, the subtherapeutic dose of an anti-tumour drug, JQ1, was loaded into CM-PEG-BPSi NPs. To illustrate ICD, we studied HMGB1 release from cell nuclei and CRT relocalization to the cell surface from CLSM images. Overall, our study aims to identify the optimal drug to synergistically induce ICD in combined cancer therapy.

**Figure 1.**
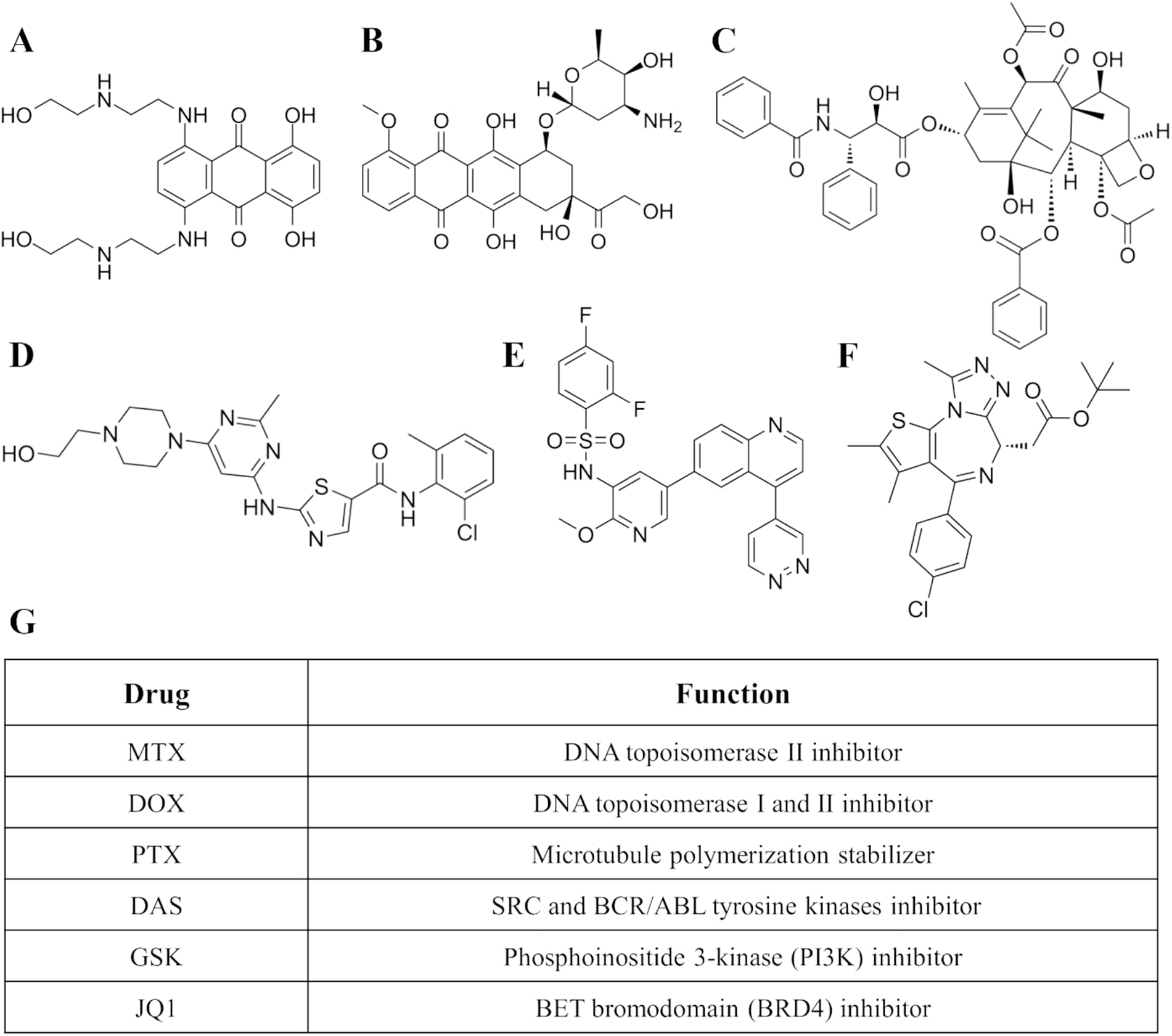
Chemical structures of (A) Mitoxantrone (MTX), (B) Doxorubicin (DOX), (C) Paclitaxel (PTX), (D) Dasatinib (DAS), (E) Omipalisib (GSK), and (F) (+)-JQ-1 (JQ1). (G) Cell growth inhibition mechanisms of selected chemotherapeutic drugs.

## 2. Materials and methods

### 2.1. Extraction of cancer cell membranes

CMs extraction was performed following the previously published method of Liu et al. (25). Briefly, MDA-MB-231 cells were cultured in RPMI (1640 w/L-Glutamine, Biowest) cell medium containing 10 % of fetal bovine serum (FBS) and 1 % of penicillin. After reaching sufficient culture disk coverage, cells were detached using trypsin and washed three times with Hanks’ Balanced Salts Solution (HBSS, Biowest) via centrifugation at 1200 rcf for 5 min. Washed cells were dispersed in hypotonic lysing buffer (20 mM of Tris-HCl pH 7.5, 10 mM of KCl, 2 mM of MgCl_2_) with one protease inhibitor cocktail tablet (Mini EDTA-free, Roche, Merck) per 10 ml. Dispersed cells were incubated with rotation at 4 °C for 0.5 h. After incubation, cells were disrupted using a Dounce homogenizer with a tight-fitting pestle and centrifuged at 3200 rcf for 5 min. Supernatants were collected. Remaining pellets were resuspended in hypotonic lysing buffer, and disruption steps were repeated. To discard pellets, collected supernatants were recentrifuged at 10 000 rcf for 10 min. To collect CM pellets, purified supernatants were centrifuged at 4 °C at 100 000 rcf for 1 h. HEPES was used to wash CM pellets via centrifugation at 100 000 rcf for 0.5 h. CM pellets were redispersed in 1 ml of HEPES. Pierce^TM^ bicinchoninic (BCA) protein assay kit (Thermo Scientific) was used to determine the CMs protein concentration by measuring the absorbance with Synergy H1 microplate reader (Biotek). Finally, CMs were stored at -20 °C for later usage.

### 2.2. Cell cytotoxicity for IC_50_ determination

Six ICD inducing drugs, MTX, DOX, PTX, DAS, GSK, and JQ1, were selected from an ICD screen of 528 oncology drugs against MDA-MB-231 cells (TNBC cell line). IC_50_ concentrations of these drugs were determined by studying their cytotoxic properties using CellTiter-Glo® 2.0 Luminescent Cell Viability Assay (Promega). MDA-MB-231 cells expressing HMGB1-GFP and CRT-RFP (stained MDA cells) were cultured in 100 µl of RPMI medium (10% FBS and 1% penicillin) in 96-well plate (PerkinElmer ViewPlate – 96 TC) at the density of 5 × 10^3^ cells/well. After 24 h incubation at 37 °C, the medium was replaced with 200 µl of RPMI containing said chemotherapeutic drugs with various concentrations. After 24 h incubation, the medium was replaced with 200 µl of fresh RPMI, continuing the incubation another 24 h. Simultaneously, the negative control was merely cultured in RPMI. Finally, the cell viability was evaluated by replacing the medium with 50 µl of both fresh RPMI and CellTiter-Glo®. The plate was incubated at room temperature for 15 min before measuring luminescent signal by Fluoroskan Ascent FL plate reader (Thermo Labsystems).

### 2.3. Preparation of BPSi NPs

(1) PEG-BPSi NPs: BPSi particles were prepared based on a novel Na-K alloy reduction method. Considering highly reactivity of the Na-K alloy (1:1, m/m), the production was performed in a glove box under argon (Ar) atmosphere. First, SiCl_4_ was mixed with toluene (0.1 g/ml). The mixture was allowed to react with the Na-K alloy in a sealed autoclave at ambient temperature for 4 h via magnetic stirring. The reaction was continued by increasing the temperature to 140 °C for another 4 h. The product was placed to a 500 °C tube oven for 2 h under N_2_ atmosphere. To quench the residual alloy, the cooled product was gradually mixed with 10 ml of acetic acid (10 wt% in hexane) allowing N2 protection and magnetic stirring at 60 °C for 4 h. Produced BPSi microparticles were purified with hydrofluoric acid (5 wt% in ethanol) before reducing particle size by ball-milling in ethanol (EtOH) at 1000 rpm for 45 min. The solution of H_2_O_2_:HCl:H_2_O (1:1:5) was used to oxidize BPSi NP surfaces at 90 °C for 15 min (26). According to the previous study of Näkki et al., generated Si-OH groups enable further NP surfaces modification with PEG-silanes (0.5 kDa and 2.0 kDa) at 110 °C for 2 h, yielding PEG-BPSi NPs as a final product (27).

(2) Cy5.5-PEG-BPSi-COOH NPs: To study homogenous cell targeting with CLSM imaging, PEG-BPSi NPs were labelled with cyanine 5.5 (NHS-Cy5.5, Lumiprobe) stain. First, 0.25 mg of NHS-Cy5.5, 10 µl of (3-Aminopropyl)triethoxysilane (APTES, Acros Organics), and 10 mg of PEG-BPSi NPs were mixed in EtOH. The mixture was stirred at 65 °C for 40 min. After stirring, obtained Cy5.5-PEG-BPSi-NH_2_ NPs were washed three times with EtOH. To turn NP surface charges from positive to negative, free NH_2_ groups on NP surfaces were capped with succinic anhydride forming COOH groups. Thus, 10 mg/ml of succinic anhydride in EtOH was mixed with 2 ml of NP solution. The mixture was rotated at room temperature overnight. Finally, obtained Cy5.5-PEG-BPSi-COOH NPs were washed three times with EtOH.

(3) CM-PEG-BPSi NPs or Cy5.5-CM-PEG-BPSi-COOH NPs: To enhance the colloidal stability and homogenous cell targeting, PEG-BPSi or Cy5.5-PEG-BPSi-COOH NPs were CM coated using wetness evaporation method. Totally 0.4 mg of NPs, 0.4 mg of CMs in HEPES, and 0.2 ml of H_2_O:PBS (1:1) were mixed via ultrasonication. Solvents were evaporated with N_2_ flow and magnetic stirring at room temperature. To remove free CMs, the sample was washed with 0.5 ml of H_2_O:PBS (1:1) by centrifuging at 10 000 rpm for 5 min. The supernatant was removed, and obtained CM-coated NPs were utilized.

(4) Drug release: To determine JQ1-loading efficiency based on the drug release, 0.5 mg of PEG-BPSi NPs, 0.5 mg of CMs, and 16.6 µg of JQ1 in DMSO were mixed via ultrasonication. According to the point (2), solvents were evaporated with N_2_ while stirring at room temperature. To remove the free drug, the sample was dispersed into 0.5 ml of H_2_O:PBS (1:1) and centrifuged at a speed of 10 000 rpm for 5 min. The supernatant was removed. To study the release kinetics of JQ1 from CM-PEG-BPSi NPs, obtained NPs were redispersed into the solution of acetonitrile and water (ACN:H_2_O, 1:1) by ultrasonication. The sample was allowed to rotate at 37 °C. At certain time periods, the sample was separated via centrifugation at 10 000 rpm for 5 min. To remove remaining impurities, supernatants were recentrifuged at 14 000 rpm for 5 min, and stored for further analyse. After each time period, remaining NPs were redispersed into 0.5 ml of fresh ACN:H_2_O (1:1). Afterwards, to estimate the released JQ1 mass, collected supernatants were analysed using high-performance liquid chromatography (HPLC) containing Inertsil® ODS-3 reversed-phase column (GL Sciences Inc., Fukushima, Japan), CBM-20A HPLC controller, CTO-10AVP column oven, LC-20AD prominence pump, LC-10ADVP pump, SIL-20AC prominence autosampler, and SPD-10AVP UV/Vis detector (Shimadzu corporation, Kyoto, Japan). The used organic mobile and aqueous phases respectively were ACN with 0.1% TFA and H_2_O with 0.1% TFA in ratio of 77:23 (v/v). The sample injection volume and the wavelength of UV/Vis detector were adjusted to 5 µl and 280 nm, respectively. Finally, the released JQ1 mass as a percentage (%) was calculated based on the following equation:

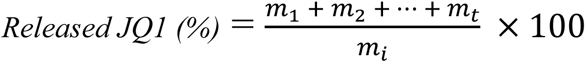

where *m_i_* is the initial mass of JQ1 and *m_t_* is the mass of JQ1 in the supernatant in each period. This released percentage was used as JQ1-loading efficiency in following experiments.

(5) JQ1-CM-PEG-BPSi NPs: JQ1-loaded CM-PEG-BPSi NPs were prepared according to the wetness evaporation method described in the point (3). As difference, JQ1 in DMSO was added into the mixed solution of NPs and CMs (1:1) based on the following equation:

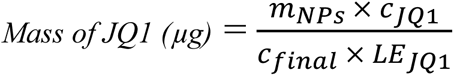

where *m_NPs_* is the mass of PEG-BPSi NPs, *c_JQ1_* is the IC_50_ concentration of JQ1, *c_final_* is the final concentration of JQ1-loaded NPs, and *LE_JQ1_* is the loading efficiency of JQ1.

The solution was evaporated with N_2_ while stirring at room temperature. To remove free CMs and JQ1, the sample was washed with 0.5 ml of H_2_O:PBS (1:1) via centrifugation at 10 000 rpm for 5 min. The supernatant was removed, and obtained JQ1-CM-PEG-BPSi NPs were utilized.

### 2.4. Physicochemical characterization

Several techniques were used to characterize OH-BPSi, PEG-BPSi, and CM-PEG-BPSi NPs. The particle size, colloidal stability, and surface charge as a zeta-potential were measured using Zetasizer Nano ZS instrument (Malvern, UK). For the measurement of NP sizes and zeta-potentials, NPs were dispersed in H_2_O at room temperature. NP colloidal stabilities were measured for 24 h, dispersing NPs in PBS (pH 7.4) at the temperature of 37 °C.

CM and PEG coatings were verified via Fourier-transform infrared spectroscopy (FTIR, Thermo Nicolet iS50). For measurements, dried 0.75 mg of calcium bromide (KBr) and 0.25 mg of NPs or pure CMs were grinded until the mixture was uniform. From mixtures, tablets were produced using Specac KBr tablet press and measured with FTIR tablet mode.

NP morphologies were studied via transmission electron microscopy (TEM, Jeol JEM-2100F). To enhance the contrast between background and CMs on NP surfaces, NPs were negative stained. First, NPs were suspended in H_2_O at 0.4 mg/ml. A drop of the suspension was placed on a carbon-coated glow copper grid (200 meshes), which was incubated 2 min at room temperature. The excess suspension was removed, and the steps were repeated. After the grid was partly dried, it was washed by a drop of H_2_O on the surface. The excess H_2_O was removed, touching the surface by a filter paper. For negative staining, 7 µl of 2% aqueous uranyl acetate stain was dropped on the grid, which was incubated 3 min. The excess stain was removed, touching the surface by a filter paper. After that, the grid was dried at room temperature.

The photothermal conversion ability of BPSi NPs was determined in photothermal heating test. NPs were dispersed in RPMI, and 200 µl of solution per well was added into a 96-well plate. The solution was heated using 808 nm laser (1 W/cm^2^) for 10 min. Three concentrations of BPSi NPs were tested: 0.025, 0.05, and 0.1 mg/ml.

### 2.5. Homotypic targeting studies

Homotypic targeting was tested using MDA-MB-231 cells while RAW264.7 cells were used as a control cell line. Cells were seeded in an 8-well plate (Ibidi) at the density of 2 × 10^4^ cells/well and cultured at 37 °C for 24 h. The medium was replaced with 50 µg/ml of Cy5.5-PEG-BPSi-COOH NPs with and without CM coating. After 4 h incubation, cells were washed three times with PBS and fixed with 4% paraformaldehyde (Sigma-Aldrich) for 10 min. Next, cells were washed twice with PBS followed by the staining with 1 µg/ml DAPI (Sigma-Aldrich) for 10 min. The staining solution was replaced with PBS. Afterwards, CLSM (Zeiss LSM 700) images with 40x magnification were taken and analysed using Interactive Microscopy Image Analysis Software (IMARIS, Oxford Instruments).

### 2.6. Photothermal therapy in vitro

PTT experiments were conducted under controlled conditions at 37 °C incubator using 3D-printed PTT system designed for 96-well plate. The system uses a proportional-integral-derivate controller to adjust an 808 nm NIR laser (A Changchun New Industries, China) output power, maintaining the targeted temperature during the heating. A MLX90621 infrared thermal array sensor (Melexis) and laser collimator were respectively used to control the temperature and adjust the laser size to fit a well. For the sensor calibration and temperature verification, K-type thermocouple was used. In all following PTT experiments, cells were heated with a power density of 1.0 W/cm^2^ for 10 min.

1. Free drug + CM-PEG-BPSi NPs + PTT: To study the potential of six chemotherapeutic drugs for ICD induction in temperature-controlled PTT, stained MDA cells were seeded in a 96-well plate (PerkinElmer ViewPlate – 96 TC) at the density of 5 ×10^3^ cells/well. Cells were cultured in 100 µl of RPMI at 37 °C for 24 h. The medium was replaced with 200 µl of free drug and CM-PEG-BPSi NP suspension in RPMI for 24 h. Free drug and NP concentrations were respectively equal to IC_50_ concentration and 0.1 mg/ml. The negative control was merely cultured in RPMI. Experiment groups were: (a) Free MTX + NPs + 45 °C laser, (b) Free DOX + NPs + 45 °C laser, (c) Free PTX + NPs + 45 °C laser, (d) Free DAS + NPs + 45 °C laser, (e) Free GSK + NPs + 45°C laser, and (f) Free JQ1 + NPs + 45 °C laser. After PTT, drug-NP suspensions were replaced with 200 µl of fresh RPMI, and cells were incubated 22 h. For CLSM imaging, cell nuclei were stained by adding 50 µl of Hoechst in RPMI (50 µg/ml) to each well. Cells were incubated at 37°C for 15 min before taking CLSM images with 20x magnification (Zeiss LSM 800 Airyscan). Finally, 24 h after PTT, cell viability was measured using CellTiter-Glo® assay.
2. CM-PEG-BPSi NPs + PTT: Stained MDA cells were cultured in 100 µl of RPMI cell medium in a 96-well plate (PerkinElmer ViewPlate – 96 TC) at the density of 5 ×10^3^ cells/well at 37 °C for 24 h. The medium was replaced with 200 µl of 0.1 mg/ml CM-PEG-BPSi NP suspension in RPMI for 24 h. The negative control was merely cultured in RPMI. Experiment groups were: (a) NPs, (b) NPs + 45 °C laser, (c) NPs + 50 °C laser, and (d) NPs + 55 °C laser. After PTT, the NP suspension was replaced with 200 µl of fresh RPMI, and cells were incubated 22 h. According to the previous point (1), cells were imaged using CLSM with 20x magnification. Finally, 24 h after PTT, cell viability was measured using CellTiter-Glo® assay.
3. JQ1-CM-PEG-BPSi NPs + PTT: Stained MDA cells were cultured in 100 µl of RPMI cell medium in a 96-well plate (PerkinElmer ViewPlate – 96 TC) at the density of 5 ×10^3^ cells/well at 37 °C for 24 h. The medium was replaced with 200 µl of 0.1 mg/ml JQ1-CM-PEG-BPSi NP suspension in RPMI for 24 h. The JQ1 concentration was 14.9 µg/ml, equal to IC_50_ concentration. The negative control was merely cultured in RPMI. Experiment groups were: (a) JQ1 NPs, (b) JQ1 NPs + 45 °C laser, (c) JQ1 NPs + 50 °C laser, and (d) JQ1 NPs + 55 °C laser. After PTT, the JQ1 NP suspension was replaced with 200 µl of fresh RPMI, and cells were incubated 22 h. According to the point (1), cells were imaged using CLSM with 20x magnification. Finally, 24 h after PTT, cell viability was measured using CellTiter-Glo® assay.

### 2.7. DAMPs analysis

The HMGB1 release from cell nuclei and CRT translocation from the ER to the cell surface were studied from CLSM images using IMARIS (Oxford Instruments). For HMGB1 release analyses, the software compares the overlap of created surfaces on cell nuclei (blue fluorescence signal) and HMGB1-GFP. As a sign of CRT translocation, the CRT-RFP intensity was analysed. To ensure accuracy, analyses were individually performed for each experiment group.

## 3. Results and discussion

### 3.1. Physicochemical characterization

BPSi particles were produced by a novel Na-K alloy reduction method. After particle size reduction, obtained BPSi NPs were oxidised to form Si-OH groups on surfaces, enabling PEGylation. Further, NPs were CM-coated and drug-loaded in wetness evaporation process.

Figure 2a shows that NP hydrodynamic diameters increased about 20-30 nm after PEGylation and CM coating, confirming their success. Moreover, both PEGylated and CM-coated NPs were stable in PBS at 37 °C for 24 h, indicating successful PEG and CM coatings (Figure 2b). In Figure 2c, zeta potentials show that OH-PEG-BPSi NPs were more negative (-31.6 mV) than PEG-BPSi NPs (-27.7 mV) and CM-PEG-BPSi NPs (-15.0 mV). Thus, especially, CM coating converted the NP surface charge towards positive, enabling higher cell targeting. FTIR spectra (Figure 2d) highlights the success of CM coating, showing transmittance peaks of proteins (1650-1550 cm^-1^) and lipids (1350-1400 cm^-1^) for pure CMs and CM-PEG-BPSi NPs. For further verification of CM coating, TEM imaging of negatively stained samples was performed. The TEM image of CM-PEG-BPSi NPs (Figure 2f) shows white external layers on NP surfaces compared to PEGylated NPs with dark clear edges (Figure 2e). This indicates a successful CM coating.

**Figure 2.**
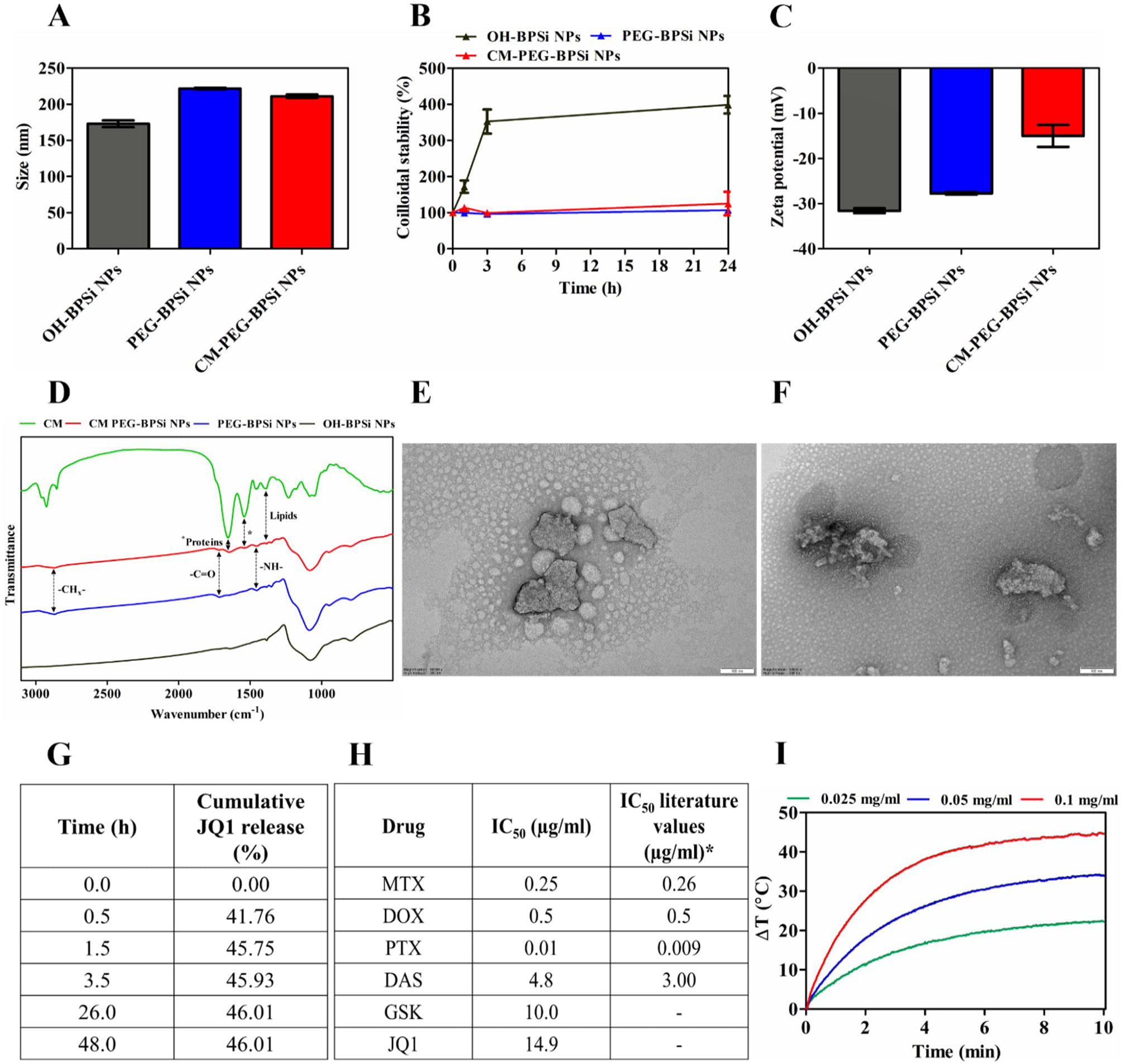
Physicochemical characterizations of NPs: **(A)** hydrodynamic diameter, **(B)** colloidal stability measured in PBS, and **(C)** zeta potential (n=3). **(D)** Fourier-transform infrared spectroscopy (FTIR) spectra of NPs and pure CMs. Transmission electron microscopy (TEM) images of negative stained **(E)** PEG-BPSi NPs and **(F)** CM-PEG-BPSi NPs. **(G)** Cumulative JQ1 release from CM-PEG-BPSi NPs measured in ACN:H_2_O (1:1). **(H)** Obtained and literature IC_50_ concentrations after 48 h incubation with MDA-MB-231 cells. (*Literature sources are presented in the section 3.1.). **(I)** Photothermal heating curves of BPSi NPs with concentrations of 0.025, 0.05, and 0.1 mg/ml.

Cumulative (%) JQ1 release from CM-PEG-BPSi NPs was studied in ANC:H_2_O (1:1) at time points 0.5, 1.5, 3.5, 26 and 48 h (Figure 2g). Most of JQ1 was already released during the first half hour. In 26 h, 46.01% of loaded JQ1 was released. After this, the release stopped. This might result from high dissolubility of JQ1 in H_2_O. However, 46.01% was applied as JQ1 loading efficiency in following experiments.

To determine IC_50_ concentrations, stained MDA-MB-231 cells were cultured with various concentrations of drugs for 24 h. After 24 h incubation, drug suspensions were replaced with fresh RPMI, and cells were incubated another 24 h before measuring cell viability. Obtained IC_50_ concentrations and corresponding literature values are showed in Figure 2h. (*Literature values: MTX (28), DOX (24), PTX (29), and DAS (30)).

The photothermal conversion ability of BPSi NPs was studied by heating NPs in RPMI in a 96-well plate with the 808 nm NIR laser (1 W/cm^2^) for 10 min. NP concentrations of 0.025, 0.05, and 0.1 mg/ml were tested. In Figure 2i, photothermal heating curves show that the highest 45 °C temperature change is obtained using 0.1 mg/ml of NPs. Thus, this concentration was selected for following PTT experiments.

### 3.2. Homotypic targeting in vitro

Homotypic targeting studies were performed by incubating MDA-MB-231 and RAW267.4 cells with Cy5.5-labelled (red fluorescence) PEG-BPSi-COOH NPs with and without CM coating for 4 h. Cell nuclei were stained with DAPI. CLSM images with 40x magnification were taken, showing the notable red fluorescence signal in the group of MDA-MB-231 cells incubated with CM-coated NPs (Figure 3a). Further, from CLSM images using IMARIS, the red fluorescence intensity on the area of cell nuclei (blue fluorescence) was analysed (Figure 3b). The analysis confirmed that in the group of MDA-MB-231 cells, Cy5.5 red fluorescence signal of CM-coated NPs was significantly higher, especially compared to the group of RAW267.4 cells with same NPs (***p ≤ 0.001). This results from CM-coated NPs recognition of cell adhesion molecules on cell surfaces, endowing successful homotypic targeting.

**Figure 3.**
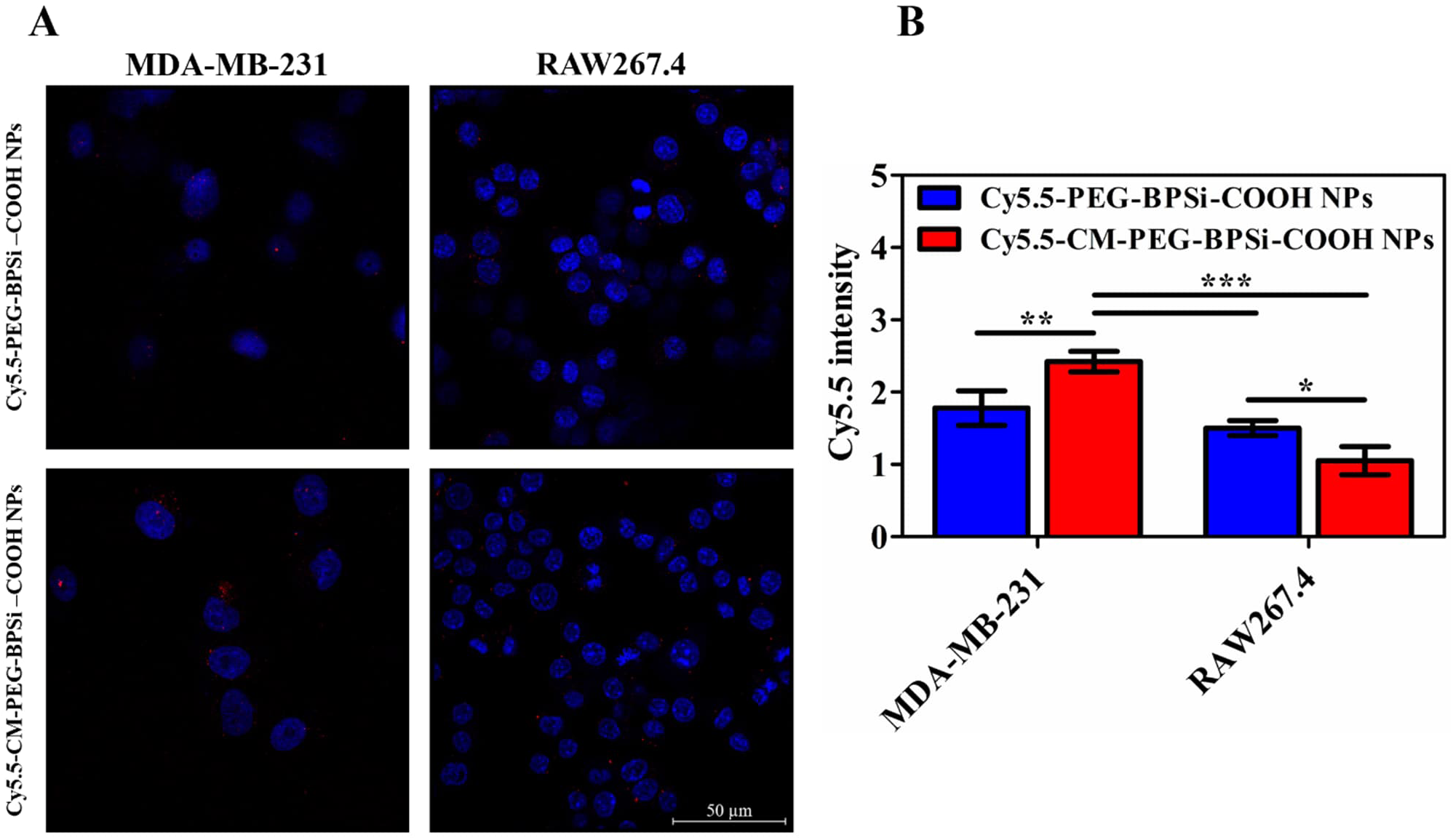
Homotypic targeting *in vitro*. **(A)** Confocal laser scanning microscopy (CLSM) images of MDA-MB-231 and RAW267.4 cells after 4 h incubation with 50 µg/ml of Cy5.5-PEG-BPSi-COOH NPs with and without CM coating. Cy5.5 (red) and DAPI (blue). Scale bar is 50 µm. **(B)** Cy5.5 fluorescence intensity on the area of cell nuclei, indicating the number of cells internalized NPs.

### 3.3. Photothermal therapy in vitro

#### 3.3.1. Free drug experiments

PTT experiments with free drugs were conducted to select the most suitable drug for ICD induction. Stained MDA-MB-231 cells were incubated with the suspension of CM-PEG-BPSi NPs and free drugs at IC_50_ concentrations for 24 h. After incubation, cells were treated using the 808 nm NIR laser (1 W/cm^2^) with 45 °C for 10 min. For DAMPs analysis, CLSM images with 20x magnification were taken, and the cell viability was measured 24 h after the treatment.

The viability result of cells treated with free drugs and PTT is showed in Figure 4a. When cells were treated with free drugs + NPs without heating, obtained viabilities predictably variated around 50 %. Merely, viabilities of MTX and DOX were around 70 %. Low cytotoxicity might result from BPSi NPs drug absorption capability, being especially sensitive for these certain molecular structures. Figure 4a shows that PTT further increases the inhibition of cancer cells growth. Compared to other groups, both free GSK and JQ1 combined with 45 °C PTT caused significant cytotoxicity (**p ≤ 0.01, ***p ≤ 0.001). However, no significant difference was observed between these groups.

**Figure 4.**
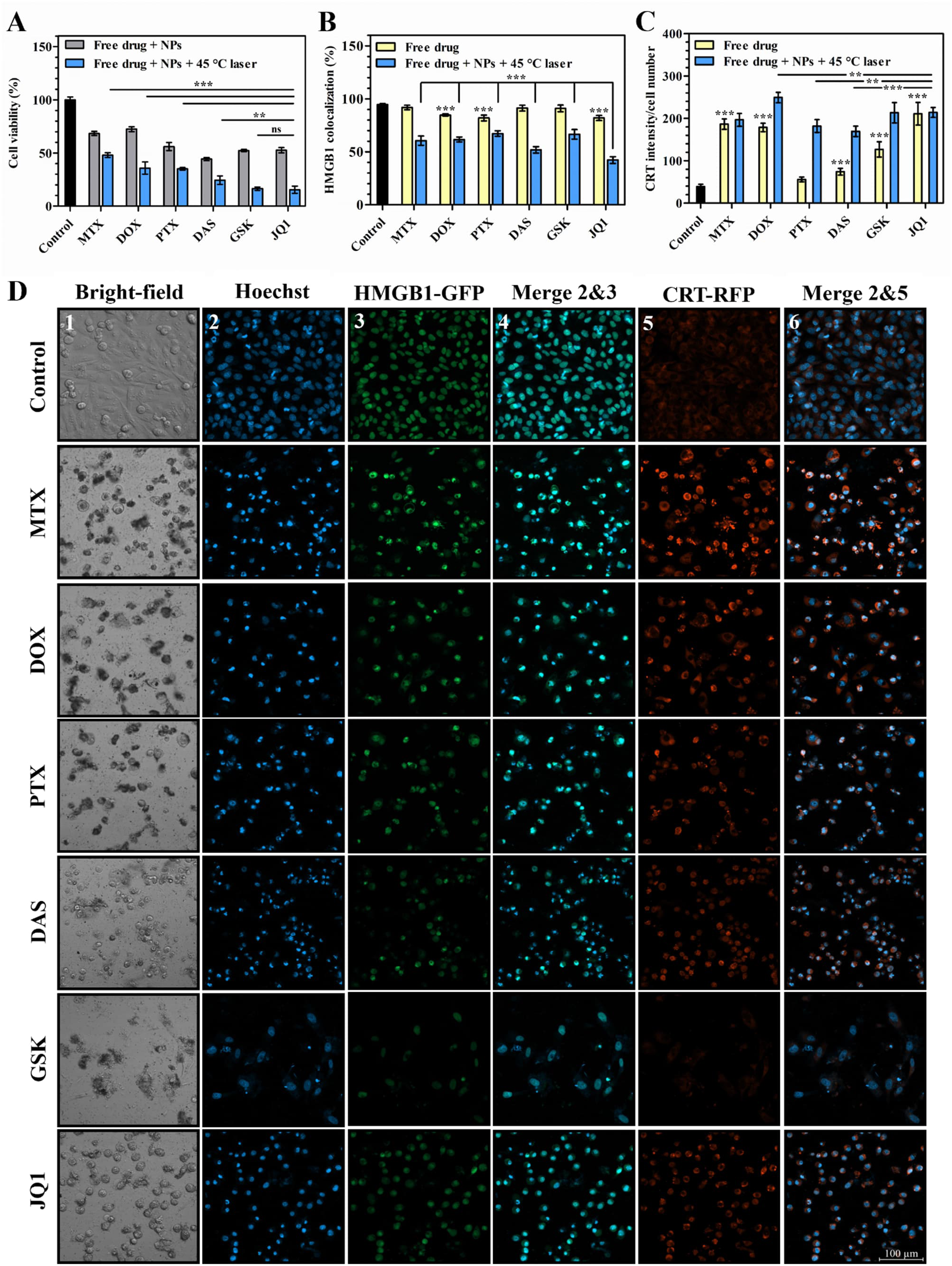
Free drug + PTT experiments. **(A)** The normalized viability of MDA-MB-231 cells treated with free drugs + CM-PEG-BPSi NPs with and without photothermal heating (45 °C laser). IC_50_ concentrations of each drug was used. The viability was measured 24 h after PTT. (n = 4). **(B)** High-mobility group box 1 (HMGB1) and cell nuclei colocalization (n = 6). **(C)** Calreticulin (CRT) intensity per cell (n = 5). **(D)** Confocal laser scanning microscopy (CLSM) images of MDA-MB-231 cells treated with free drugs and 45 °C PTT. Cell nuclei were stained with Hoechst (blue). Scale bar is 100 µm. Significance: **p ≤ 0.01 and ***p ≤ 0.001.

As markers of ICD, HMGB1 cytoplasmic release and CRT relocalization from the ER to the cell surface were analysed from CLSM images (Figure 4d). Figure 4b shows the overlapping area of HMGB1-GFP and cell nuclei (Hoechst) as HMGB1 colocalization. Compared to control group, free DOX, PTX, and JQ1 significantly decreased the colocalization (***p ≤ 0.001). This is consistent with previous studies, which have demonstrated the effectiveness of these drugs to promote HMGB1 release (31–33). When free drugs were combined with 45 °C PTT, the colocalization noticeably decreased. However, JQ1 caused the most significant decrease in the HMGB1 colocalization (***p ≤ 0.001). This might be due to the mechanism of JQ1 to cause cellular stress reactions, activating specific signalling pathways to release HMGB1 via ICD (34).

In ICD, CRT translocates from the ER to the cell surface, binding to certain membrane receptors (7). Simultaneously, it aggregates. This phenomenon can be seen as increased CRT-RFP intensity (Figure 4d). Figure 4c shows CRT intensity per cell number. Except PTX, free drugs significantly increased CRT intensity compared to the control group (***p ≤ 0.001). After PTT, the increase in CRT intensity varied a lot. For PTX, DAS, and GSK, the increase was notable, but for the other groups, the change was minor. However, free DOX combined with 45 °C PTT showed the significantly highest CRT intensity (**p ≤ 0.01). In addition, the intensity was high in GSK and JQ1 groups. According to the JQ1 group, PTT was ineffective on CRT intensity. This might be the result of similar mechanism of PTT and JQ1 to cause CRT translocation via apoptotic cell death (35,36). In addition, both could promote CRT translocation by inducing the ER stress (37).

In summary, free drug PTT experiments showed that among the drugs, GSK and JQ1 are the most effective to kill the cancer cells (Figure 4a). Free JQ1 combined with 45 °C PTT significantly decreased the HMGB1 colocalization (Figure 4b). The highest CRT intensity was achieved when free DOX was combined with 45 °C PTT (Figure 4c). Thus, based on these results, JQ1 indicated to be the most suitable drug to induce ICD in PTT, and was selected for the following PTT experiments.

#### 3.3.2. PTT experiments with drug-loaded NPs

Based on free drug experiments, JQ1 was selected as the most effective drug to induce ICD. Next, different PTT temperatures were tested to evaluate the optimal temperature for ICD induction. Shortly, stained MDA-MB-231 cells were incubated with JQ1-loaded CM-PEG-BPSi NPs for 24 h. JQ1 concentration was equal to the IC_50_ value (Figure 2h). After incubation, cells were treated using the 808 nm NIR laser (1 W/cm^2^) with temperatures of 45 °C, 50 °C, and 55 °C for 10 min. CM-PEG-BPSi NPs were used as control groups to evaluate the effect of PTT. For DAMPs analysis, CLSM images with 20x magnification were taken, and the cell viability was measured 24 h after the treatment.

Figure 5a shows the viability of cells treated with PTT using JQ1- and CM-PEG-BPSi NPs. First, in the cell viability, temperature-dependent relationship was observed. The cell viability against CM NPs without heating was equal to the control group. Thus, with the concentration of 0.1 mg/ml, CM NPs were noncytotoxic. With JQ1 NPs without heating, the cell viability was predictably around 50 %. This indicates the successful JQ1-loading into and -release from NPs. In the group of 45 °C PTT, significant difference in the cytotoxicity between CM and JQ1 NPs was observed (***p ≤ 0.001). Combined therapy of JQ1 and 45 °C PTT inhibited the growth of cancer cells by 75 %. Compared to pure 50 °C PTT, the result was equal. Combining JQ1 with 50 °C PTT, the cytotoxicity increased by half (***p ≤ 0.001). Moreover, between groups of 50 °C and 55 °C PTT, significant difference was observed for both CM and JQ1 NPs (***p ≤ 0.001 and *p ≤ 0.05, respectively). In the group of 55 °C PTT, the difference between CM and JQ1 NPs was insignificant. This is due to the dominance of necrosis in cell death caused by the high temperature (35).

**Figure 5.**
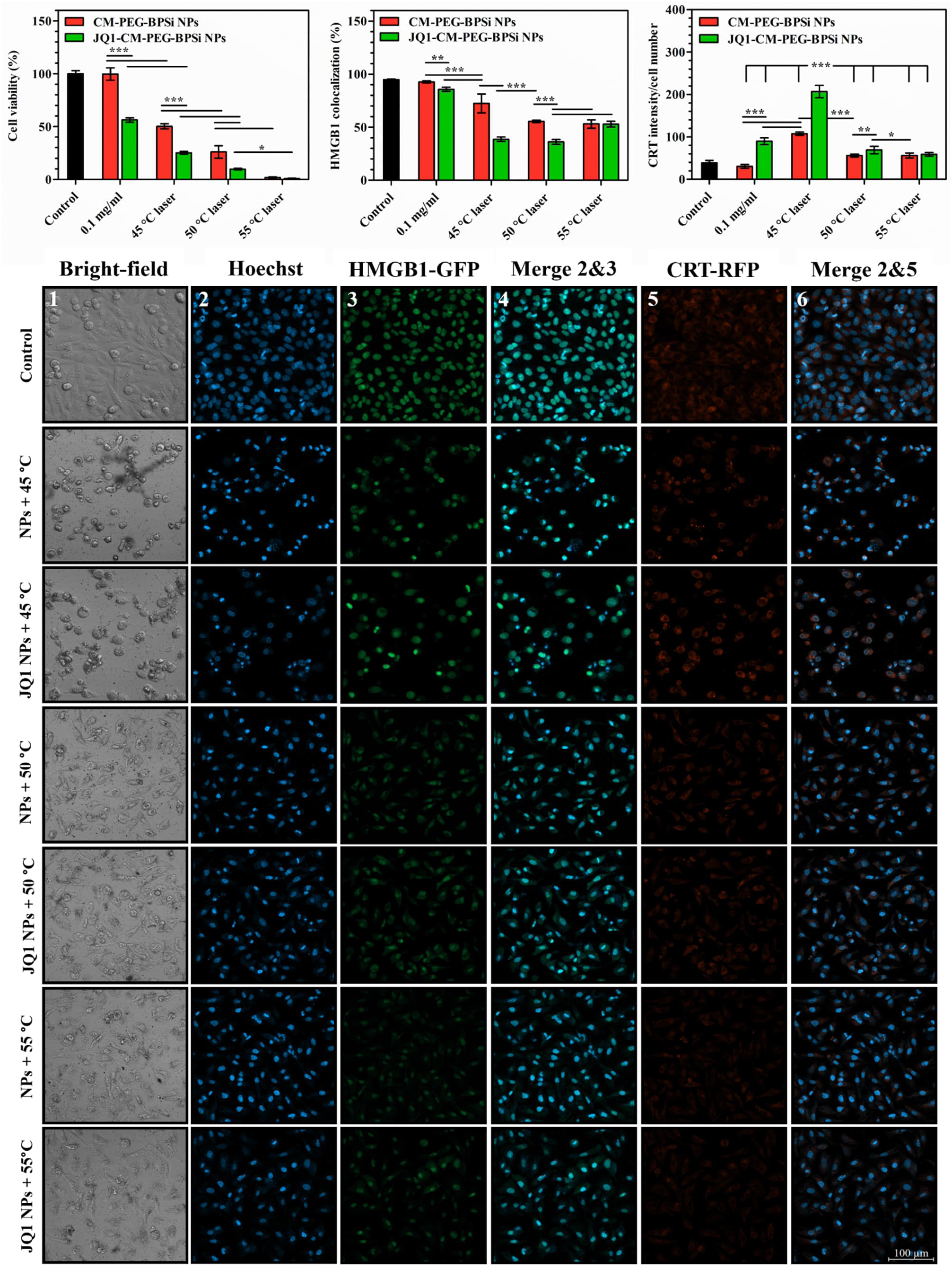
PTT experiments with JQ1-loaded NPs. **(A)** The normalized viability of MDA-MB-231 cells treated with JQ1- and CM-PEG-BPSi NPs with and without photothermal heating. Temperatures of 45 °C, 50 °C, and 55 °C were used. JQ1 concentration was equal to the IC_50_ value. The viability was measured 24 h after PTT. (n = 4). **(B)** High-mobility group box 1 (HMGB1) and cell nuclei colocalization (n = 6). **(C)** Calreticulin (CRT) intensity per cell (n = 5). **(D)** Confocal laser scanning microscopy (CLSM) images of MDA-MB-231 cells treated with 45°C, 50 °C, and 55 °C PTT using JQ1- and CM-PEG-BPSi NPs. Cell nuclei were stained with Hoechst (blue). Scale bar is 100 µm. Significance: *p ≤ 0.05, **p ≤ 0.01, and ***p ≤ 0.001.

As in the section of 3.3.1., markers of ICD were analysed from CLSM images (Figure 5d). Figure 5b shows the HMGB1 colocalization result, when JQ1- and CM-PEG-BPSi NPs were applied in PTT. The HMGB1 colocalization caused by CM NPs without heating is unsignificant compared to the control group. This indicates that NPs has no disturbing effect on cell functions. Compared to JQ1 NPs without heating, significant difference was observed (**p ≤ 0.01). The result is equal with free JQ1 (Figure 4b), indicating successful JQ1-loading and -release from NPs. In the group of 45 °C PTT, the difference between CM and JQ1 NPs was significant (***p ≤ 0.001). 45 °C PTT with JQ1 NPs decreased HMGB1 colocalization even more than pure 50 °C PTT (***p ≤ 0.001). However, compared to 50 °C PTT with JQ1 NPs, no significance was observed. The HMGB1 colocalization again increased when 55 °C PTT was applied. Among the group, no significance between CM and JQ1 NPs was observed. Thus, with high temperatures, JQ1 has no effect on HMGB1 release. This indicates that cells were mainly killed via necrosis, suddenly releasing HMGB1.

CRT intensity per cell number was analysed from CLSM images (Figure 5c). Within all the groups, 45 °C PTT with JQ1-loaded NPs showed the highest CRT intensity (***p ≤ 0.001). In addition, considering the groups of CM NPs, the temperature of 45 °C achieved the highest CRT intensity (***p ≤ 0.001). 45 °C PTT is well-known to cause mild hyperthermia in cells, leading to apoptotic cell death. Subsequently, “eat me” signals are produced by cells undergoing apoptosis (38). One of these signals might be observed as presence of CRT on the cell surface. This is further promoted by JQ1, which inhibits bromodomain-containing protein 4 (BRD4). Via BRD4 inhibition, JQ1 can affect transcriptional regulation, enabling intracellular stress reactions. Further, BRD4 inhibition enhances transcriptional factor NF-ĸB activation, increasing inflammatory cytokines production (39). Moreover, these cytokines could induce the ER stress, and thus strong CRT translocation (40,41).

Compared to the control group, 0.1 mg/ml of CM NPs have no significant effect for CRT intensity (Figure 5c). JQ1 NPs without heating increased the CRT intensity, indicating JQ1 release from NPs. In the group of 50 °C PTT, significant difference between CM and JQ1 NPs were observed (**p ≤ 0.01). However, comparing CRT intensities between CM and JQ1 NPs in the groups of 50 °C and 55 °C PTT, respectively, no significant difference was observed. This results from the dominance of high temperatures over the drug. In PTT, high temperatures cause necrotic cell death, in which plasma membrane (outer part of cell membrane) is damaged, rapidly releasing DAMPs into extracellular phase (38). Thus, CRT translocation on the cell surface might be prevented.

## 4. Conclusion

The present study utilized CM-coated BPSi NPs as both drug carriers and PCAs to demonstrate the efficiency of combined CHT-PTT in inducing ICD. The results confirmed successful CMs coating of NPs, enabling their good colloidal stability and homotypic targeting properties. Six chemotherapeutic drugs with different cell growth inhibition mechanisms were tested to compare their ICD induction potential. It was found that BRD4 protein inhibitor, JQ1, was the most effective among these drugs after analysing ICD induction from CLSM images. Finally, we demonstrated JQ1-loaded NPs at mild hyperthermia (45 °C) was the best setting combined CHT-PTT in ICD induction. Overall, this study provides a rational design of CHT-PTT for efficient cancer immunotherapy.

## Acknowledgements

This study was supported by the funding from Research Council of Finland (Grant no. 356056, 359706, and 356992). Cell and Tissue Imaging Unit (UEF) and SIB Labs at UEF were acknowledged for technical support.

